# Contralateral Prefrontal and Network Engagement during Left DLPFC 10 Hz rTMS: An Interleaved TMS-fMRI Study in Healthy Adults

**DOI:** 10.1101/2025.04.14.648588

**Authors:** Timo van Hattem, Kai-Yen Chang, Martin Tik, Paul Taylor, Jonas Björklund, Lucia Bulubas, Frank Padberg, Daniel Keeser, Mattia Campana

**Affiliations:** Department of Psychiatry and Psychotherapy, University Hospital LMU, LMU Munich, Germany; Neuroimaging Core Unit Munich - NICUM, University Hospital LMU, LMU Munich, Germany; Hertie-Institute for Clinical Brain Research, University of Tübingen, Germany; Department of Neurology & Stroke, University of Tübingen, Germany; High Field MR Center, Center for Medical Physics and Biomedical Engineering, Medical University of Vienna, Vienna, Austria; Brain Stimulation Lab, Department of Psychiatry and Behavioral Sciences, Stanford University, Stanford, USA; Department of Psychology, LMU Munich, Germany; Department of Psychology, Universität Zürich, Switzerland; DZPG (German Center for Mental Health), partner site Munich-Augsburg, Germany; Munich Center for Neurosciences (MCN), Ludwig Maximilian University LMU, Munich, Germany; LVR Hospital, Department of General Psychiatry 2, Clinic of the Heinrich Heine University, Düsseldorf, Germany

**Keywords:** Interleaved TMS-fMRI, 10 Hz rTMS, iTBS, major depression, neuromodulation, neuroimaging

## Abstract

**Background:** High-frequency repetitive transcranial magnetic stimulation (rTMS) over the left dorsolateral prefrontal cortex (DLPFC) serves as an effective treatment for major depression and other psychiatric disorders. Despite its growing clinical application, the neural mechanisms by which prefrontal rTMS exerts its therapeutic effects remain incompletely understood. To address this gap, we investigated the immediate blood-oxygen-level-dependent (BOLD) activity during 600 stimuli of left DLPFC 10 Hz rTMS in healthy individuals using interleaved TMS-fMRI.

**Methods:** In a crossover design, 17 healthy subjects received 10 Hz rTMS (60 trains with 9-second intertrain intervals) over the left DLPFC at 40% and 80% of their resting motor threshold (rMT) inside the MR scanner.

**Results:** 10 Hz rTMS over the left DLPFC elicited BOLD responses in prefrontal regions, cingulate cortex, insula, striatum, thalamus, as well as auditory and somatosensory areas. Notably, our findings revealed that 10 Hz rTMS effects were lateralized towards the contralateral (right) DLPFC. Dose-response effects (40% vs. 80% rTMS) were exclusively observed in contralateral subcortical regions, whereas dose-responses within the DLPFC showed substantial inter-individual variability.

**Conclusions:** The 10 Hz rTMS protocol used in this study induced distinct target engagement and propagation patterns in the prefrontal cortex. These patterns differ from our previous interleaved TMS-fMRI findings using 600 stimuli of left DLPFC intermittent theta burst stimulation (iTBS) at the same intensities. Thus, interleaved TMS-fMRI emerges as a valuable method for comparing clinical prefrontal rTMS protocols regarding their immediate effect on brain circuits in order to differentiate their action mechanisms and to potentially inform clinical applications.

## 1. Introduction

Repetitive transcranial magnetic stimulation (rTMS) has emerged as an effective therapeutic intervention for various psychiatric and neurological disorders (Hyde et al., 2022; Lisanby et al., 2024). rTMS can modulate pathological excitability and connectivity in brain networks, resulting in symptom relief that extends beyond the immediate period of stimulation (Klomjai et al., 2015; Lefaucheur et al., 2020). High-frequency 10 Hz rTMS over the left dorsolateral prefrontal cortex (DLPFC) has become a FDA-approved clinical standard for treating pharmacoresistant major depressive disorder (MDD) (O’Reardon et al., 2007; Berlim et al., 2014; McClintock et al., 2018; Papakostas et al., 2024).

While the therapeutic benefits of rTMS are well-established (George et al., 2013; Lefaucheur et al., 2020; Kan et al., 2023), key questions persist about how rTMS engages target regions and propagates its effects through (sub)cortical networks (Siebner et al., 2022). Local and remote effects of rTMS vary depending on protocol parameters, such as stimulation frequency and intensity, with different approaches potentially recruiting distinct interneuronal circuits (Di Lazzaro et al., 2011; Hamada et al., 2013). Understanding the immediate changes in neural activity induced by rTMS across the entire brain can provide crucial insights into their underlying neurophysiological mechanisms (Tik et al., 2023a, 2023b; Chang et al., 2024a, 2024b).

Combining TMS with functional magnetic resonance imaging (fMRI) allows investigating acute effects on cerebral activation and connectivity with high spatial accuracy (Bohning et al., 1998; Bergmann et al., 2021; Mizutani-Tiebel et al., 2022). Previous TMS-fMRI research in healthy people has primarily used single-pulse TMS or short 10 Hz bursts on the primary motor cortex (M1) and DLPFC (Bergmann et al., 2021; Mizutani-Tiebel et al., 2022; Xia et al., 2024). Investigating stimulation protocols inside the MR scanner that are closer to rTMS protocols in therapeutic applications could enhance the translational value of interleaved TMS-fMRI techniques by demonstrating target involvement and potentially helping identify predictive biomarkers of treatment response.

In this study, we investigate the immediate blood-oxygen-level-dependent (BOLD) responses to a 600-stimuli 10 Hz rTMS protocol at two stimulation intensities (i.e. 40% and 80% resting motor threshold, rMT) targeting the left DLPFC in healthy, resting individuals. This parameter selection was guided by our previous study, which used an identical interleaved TMS-fMRI setup and a similar crossover design to examine the cortical effects of a full 600-stimuli intermittent theta burst stimulation (iTBS) protocol (Chang et al., 2024a, 2024b). Accordingly, we also aimed to provide a descriptive comparison of the direct neural effects of 10 Hz rTMS and iTBS under matched stimulation parameters. The current study on 10 Hz rTMS, alongside the narrative comparison of its findings with iTBS, may help deepen our mechanistic understanding of how high-frequency rTMS acutely modulate local and remote neural activity, which could provide foundational knowledge to improve future neuromodulation treatments.

## 2. Methods and Materials

### 2.1 Participants

Twenty healthy subjects (13 females; mean age = 29.25 ± 7.77 years) were recruited for this study. Participants had no contraindications to TMS and MRI, and no history of psychiatric or neurological disorders. All participants provided written informed consent before the experiment. The study was approved by the ethical committee of LMU Munich and was conducted in accordance with guidelines of the Declaration of Helsinki.

Two participants dropped out during the study due to mild adverse events associated with active TMS (i.e., migraine after TMS, intolerable pressure of the TMS coil on the scalp) and one participant was excluded due to excessive motion, resulting in a total of 17 subjects (11 females; mean age = 28.18 ± 8.02 years) included for the final data analyses (Table S1).

### 2.2 Experimental design

This study comprised a baseline measurement and two experimental sessions with interleaved TMS-fMRI (Figure 1A). During the baseline session, subjects underwent structural MRI for neuronavigation. Individual rMT was determined for each subject while lying supine inside the MR environment. rMT was defined as the intensity that evoked a motor-evoked potential (MEP) with a peak-to-peak amplitude greater than 50 µV in five out of ten trials (Rossini et al., 1994). The average rMT was 74% (SD = 9%) expressed as percentage of maximal stimulator output (MSO) (Table S1). During the second and third sessions, subjects received 10 Hz rTMS over the left DLPFC inside the MR scanner with a stimulation intensity of either 40% rMT or 80% rMT. The intensity was randomized across sessions and the sessions were separated by a minimum of one week.

**Figure 1.**
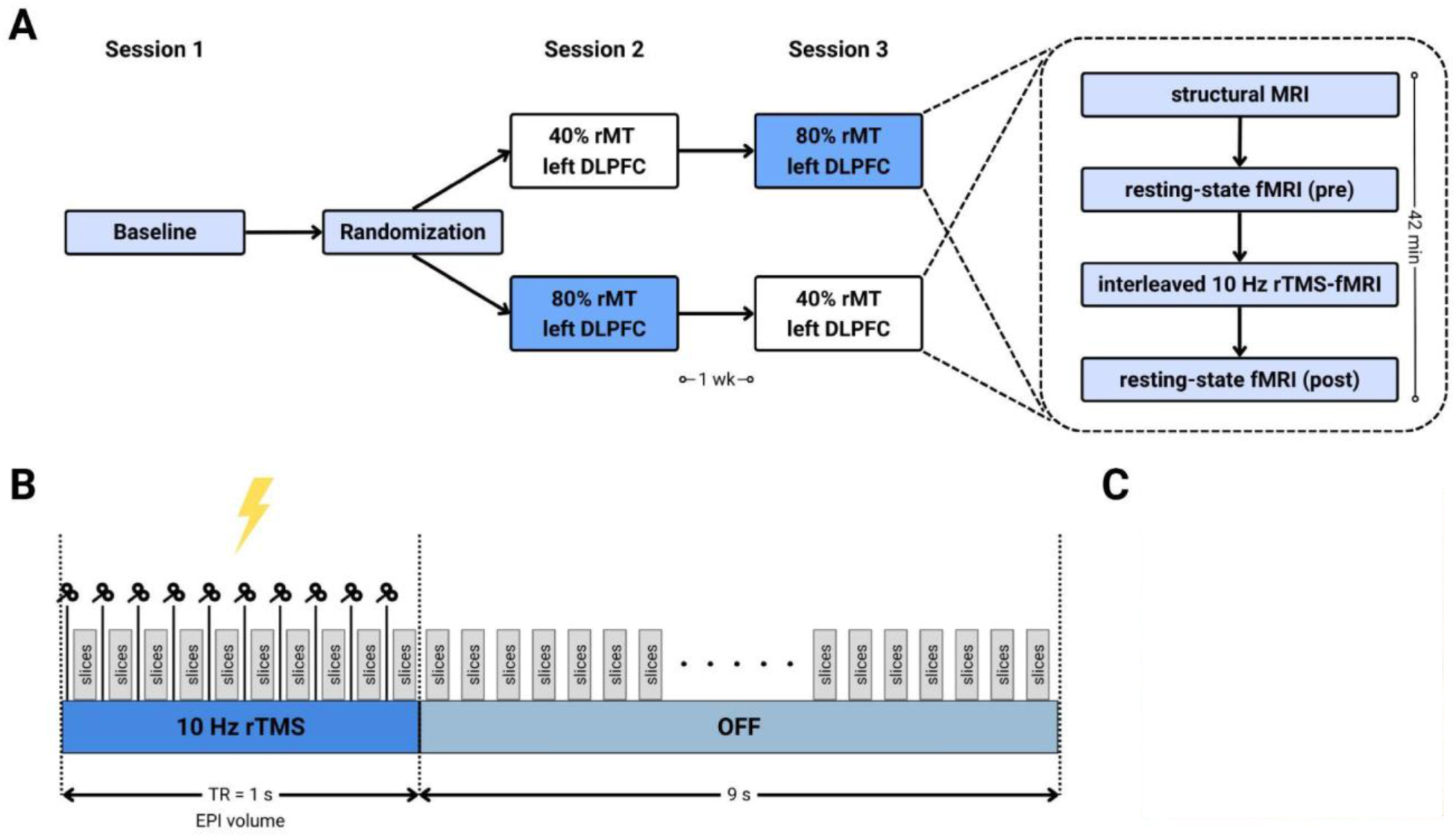
Experimental design, stimulation protocol, and setup of interleaved 10 Hz rTMS-fMRI experiment. (A) Randomized, crossover study design with a baseline measurement and two interleaved TMS-fMRI sessions. Each experimental session included structural MRI, pre- and post-resting-state fMRI, and interleaved 10 Hz rTMS-fMRI over the left DLPFC at either 40% or 80% of the resting motor threshold (rMT). (B) TMS pulses in 1-second trains at 10 Hz were interleaved with multi-band EPI slices, followed by 9 seconds of rest. (C) Interleaved TMS-fMRI setup with two 7-channel surface RF coils placed on each side of the head to cover the entire brain.

### 2.3 TMS

TMS was delivered using a MagPro X100 stimulator and a MR-compatible MRi-B91 TMS coil (MagVenture A/S, Farum, Denmark). TMS was applied over the left DLPFC (x, y, z = - 38, 44, 26) (Blumberger et al., 2018) using neuronavigation (Localite GmbH, Bonn, Germany) (Figure 1C). The 10 Hz rTMS protocol consisted of 60 trains with 10 pulses per train (i.e., interpulse interval of 0.1 seconds) and a 9-second intertrain interval, totaling 600 pulses per TMS-fMRI session (Figure 1B). TMS pulses were interleaved with multi-band EPI slices. The current protocol is a modification from the standard clinical application of 10 Hz rTMS, which typically involves 3000 pulses (4 s ON, 26 s OFF) per session over 37.5 minutes (O’Reardon et al., 2007; Blumberger et al., 2018). Note that we reduced the number of TMS pulses to allow a direct comparison with a full iTBS protocol (i.e., 600 pulses/session) (Huang et al., 2005), such as from our previous TMS-fMRI study (Chang et al., 2024b).

### 2.4 Image acquisition

Imaging data were acquired using a 3T Siemens PRISMA MRI-scanner (Siemens, Erlangen, Germany). T1-weighted anatomical images for neuronavigation were acquired during the baseline session using a standard 64-channel head/neck coil (TE = 2.26 ms, TR = 2300 ms, TA = 5:21 m, TI = 900 ms, flip angle = 8°, voxel size = 1.0 × 1.0 × 1.0 mm, number of slices = 192, slice thickness = 1 mm, FOV = 256 mm). During the interleaved TMS-fMRI sessions, two 7-channel surface RF coils were placed on each side of the front of the head to cover the entire brain (Figure 1C) (Navarro de Lara et al., 2015). Structural images were acquired using a magnetization-prepared rapid gradient-echo (MP2RAGE) sequence (TE = 2.98 ms, TR = 4000 ms, TA = 6:26 m, TI = 700 ms, flip angle = 4°, voxel size = 1.0 × 1.0 × 1.0 mm, number of slices = 160, slice thickness = 1 mm, FOV = 256 mm). Multi-band accelerated echo planar imaging (EPI) sequences were used for interleaved TMS-fMRI (TE = 38 ms, TR = 1000 ms, TA = 10:27 m, flip angle = 60°, voxel size = 3.0 × 3.0 × 3.0 mm, number of slices = 40, slice thickness = 3 mm, FOV = 192 mm, MB-factor = 4).

### 2.5 Data analysis

#### 2.5.1 Preprocessing

fMRI data were preprocessed using the methods described in Chang et al. (2024a, 2024b). In brief, anatomical images were segmented and normalized to Montreal Neurological Institute (MNI) standard space. Functional images underwent bias-field correction, despiking, motion correction, coregistration with anatomical images, normalization, and spatial smoothing. Subjects with a mean framewise displacement greater than 0.3 mm were excluded from all further analysis (Table S1) (Power et al., 2012). An independent component analysis (ICA) was performed to reduce physiological noise (e.g., motion, cerebrospinal fluid (CSF) pulsations) and artifacts that may have been introduced by the interleaved TMS-fMRI setup (e.g, leakage currents, RF interference due to the TMS hardware) (Griffanti et al., 2017; Bergmann et al., 2021; Mizutani-Tiebel et al., 2022; Riddle et al., 2022). Data quality was checked after each preprocessing step via visual inspection. For more details, see Supplementary Materials.

#### 2.5.2 Whole-brain BOLD-fMRI

Whole-brain analysis was conducted on the denoised data in SPM12 (http://www.fil.ion.ucl.ac.uk/spm/software/spm12/) to test for brain-wide BOLD signal changes in response to 10 Hz rTMS. At the single-subject level, a linear regression was performed for each voxel using a generalized least squares method and a global approximate AR(1) autocorrelation model. A high-pass filter (128 s cut off = 0.008 Hz) using discrete cosine transform basis sets was used to remove high-frequency noise components. The regressor of interest modeled the blocked periods of stimulation at 10 Hz and was convolved with a canonical hemodynamic response function (HRF). Realignment parameters were included in the model as nuisance regressors. Individual subject-level parameter estimates (beta weights, a.u.) from the model were extracted to compute group-level averages. Statistical significance was tested against an implicit baseline with one-sample t-tests applying a p < .001 voxel-level threshold and a p < .05 Family-Wise Error (FWE) cluster-level threshold.

#### 2.5.3 Regions-of-interest

A region-of-interest (ROI) analysis was conducted to compare evoked BOLD responses across stimulation intensities. A spherical ROI (radius = 10 mm) was created for the left DLPFC centered on the stimulation target (x, y, z = -38, 44, 26) (Blumberger et al., 2018; Chang et al., 2024a, 2024b) and for the right DLPFC on the contralateral homologous location (x, y, z = 38, 44, 26) using the MarsBaR toolbox for SPM (Brett et al., 2002). Additional spherical ROIs were generated for bilateral anterior insula, rostral ACC, and dorsal caudate nucleus. Their sizes were modified where necessary to prevent anatomical overlap. These ROIs were selected *a priori* based on previous work indicating that left DLPFC stimulation activates interconnected salience and striatal networks (Hanlon et al., 2013; Hawco et al., 2018; Riddle et al., 2022). See Table S2 and Figure S1 for sphere radius and MNI coordinates of center of mass for each ROI.

Two-way repeated measures ANOVA were performed to test for main effects of (and interaction between) intensity of stimulation and hemisphere on BOLD responses. Post-hoc pairwise comparisons of mean activations in ROIs were conducted using two-tailed paired t-tests (p < .05, FWE Holm-Bonferroni correction).

## 3. Results

### 3.1 Interleaved 10 Hz rTMS-fMRI

A whole-brain analysis assessed BOLD activation evoked by 40% rMT and 80% rMT 10 Hz rTMS over the left DLPFC. Stimulation at 40% rMT increased BOLD signal in the right middle frontal gyrus, superior frontal gyrus, and supramarginal gyrus, as well as bilateral insula, thalamus, and striatum (Figure 2A, Table 1). No significant BOLD activation was observed at the stimulation site in the left DLPFC at 40% rMT. At 80% rMT, 10 Hz rTMS evoked activity in both the left and right middle frontal gyri (Figure 2B, Table 1). Additionally, widespread BOLD activation was found in remote cortical areas, including bilateral insula, ACC, precentral gyrus, supramarginal gyrus, middle temporal gyrus, superior parietal gyrus, thalamus, and striatum. Notably, 80% rMT stimulation evoked a stronger BOLD signal increase in the contralateral (right) hemisphere than the hemisphere stimulated. A significant difference between the two intensities on the whole-brain level was only found in the hippocampus (Figure S2). There was no correlation between simulated E-field strength and BOLD responses in the left DLPFC (Figure S3).

**Figure 2.**
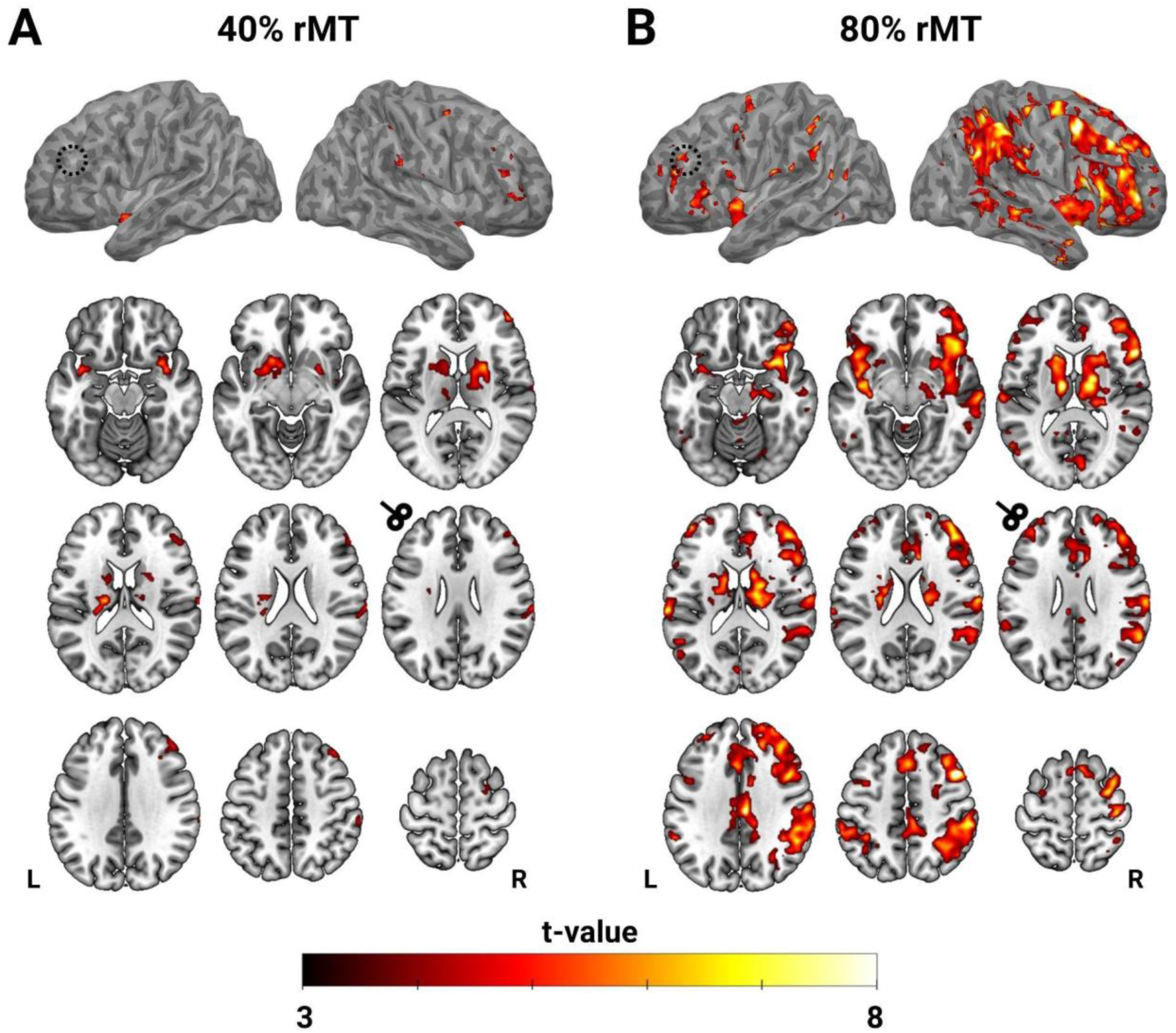
Group-level activation maps of 10 Hz rTMS-evoked BOLD responses at 40% rMT (A) and 80% rMT (B). All activation maps are thresholded at voxel-level p < .001 and cluster-level p < .05 FWE corrected. Axial slices display MNI Z-coordinates: -16, -8, 12; 18, 22, 26; 34, 48, 62. Dashed circle and TMS-coil pictogram roughly indicate target location.

**Table 1.** Peak BOLD activation in significant clusters during 10 Hz rTMS at 40% rMT and 80% rMT. Results are thresholded at voxel-level p < .001 and cluster-level p < .05 FWE corrected.

| Region | Hemisphere | Peak MNI coordinates (x, y, z) | t-value | z-value |
| --- | --- | --- | --- | --- |
| **40% rMT** |  |  |  |  |
| Middle frontal gyrus | R | 46, 52, 12 | 6.73 | 4.57 |
|  |  | 44, 38, 38 | 6.12 | 4.33 |
| Superior frontal gyrus | R | 24, -4, 56 | 6.17 | 4.35 |
| Insula | L | -40, 4, -14 | 6.05 | 4.30 |
|  | R | 34, 10, -16 | 6.07 | 4.31 |
| Supramarginal gyrus | R | 68, -26, 28 | 5.46 | 4.04 |
|  |  | 58, -32, 54 | 5.87 | 4.23 |
| Striatum | L | -20, 16, -2 | 6.88 | 4.63 |
|  | R | 24, 4, 14 | 7.71 | 4.91 |
| Thalamus | L | -18, -18, 18 | 6.55 | 4.50 |
|  | R | 12, -4, 6 | 6.62 | 4.53 |
| **80% rMT** |  |  |  |  |
| Middle frontal gyrus | L | -44, 48, 18 | 7.15 | 4.73 |
|  | R | 46, 14, 48 | 11.42 | 5.88 |
|  |  | 42, 52, 2 | 9.24 | 5.36 |
| Superior frontal gyrus | L | -8, 10, 58 | 6.55 | 4.50 |
|  |  | -26, -6, 58 | 5.07 | 3.86 |
|  | R | 12, 10, 70 | 6.70 | 4.56 |
| Inferior frontal gyrus | L | -52, 28, 4 | 5.59 | 4.10 |
|  | R | 56, 20, 10 | 10.12 | 5.59 |
| Anterior/middle cingulate gyrus | L | -2, 20, 36 | 6.42 | 4.45 |
|  | R | 10, 36, 18 | 5.30 | 3.97 |
| Posterior cingulate gyrus | L | -4, -30, 42 | 5.85 | 4.22 |
|  | R | 6, -22, 34 | 9.07 | 5.32 |
| Insula | L | -42, 2, -10 | 6.63 | 4.54 |
|  | R | 32, 10, -16 | 8.05 | 5.02 |
| Precentral gyrus | R | 42, -20, 60 | 7.32 | 4.78 |
|  |  | 32, -4, 60 | 7.53 | 4.86 |
| Supramarginal gyrus | L | -60, -42, 46 | 6.69 | 4.56 |
|  | R | 58, -32, 46 | 8.03 | 5.01 |
| Middle temporal gyrus | R | 58, -28, -4 | 6.77 | 4.59 |
| Superior temporal gyrus | L | -64, -26, 16 | 7.84 | 4.96 |
| Superior parietal lobule | L | -38, -46, 50 | 6.57 | 4.51 |
|  | R | 38, -48, 56 | 8.77 | 5.23 |
| Striatum | L | -16, 14, 14 | 7.03 | 4.68 |
|  | R | 14, 4, 20 | 6.63 | 4.53 |
| Thalamus | L | -10, -12, 12 | 8.87 | 5.26 |
|  | R | 14, -12, 14 | 8.85 | 5.26 |

### 3.2 ROI analyses

A subsequent series of ROI analyses was conducted to directly compare BOLD responses evoked by 10 Hz rTMS across different stimulation intensities and to verify the observed lateralization in neural activity. We observed a significant main effect of stimulation intensity bilaterally in the insula [F(1,16) = 10.182, p = 0.006], but not in the DLPFC [F(1,16) = 3.194, p = 0.093], nor in the ACC [F(1,16) = 3.653, p = 0.074], nor in the caudate [F(1,16) = 0.735, p = 0.404]. Specifically, BOLD activation was significantly greater at 80% rMT than 40% rMT in both the left insula [t(16) = -3.01, p = 0.032] and right insula [t(16) = -2.87, p = 0.033] (Figure 3B). Additionally, there was a significant main effect of the hemisphere in the DLPFC [F(1,16) = 15.337, p = 0.001] and insula [F(1,16) = 6.518, p = 0.021]. Post-hoc analysis revealed that BOLD signals were significantly greater in the right DLPFC than in the left DLPFC at both 40% rMT [t(16) = -2.98, p = 0.027] and 80% rMT [t(16) = -3.67, p = 0.008] (Figure 3A), and in the insula at 80% rMT [t(16) = -2.50, p = 0.048] (Figure 3B). Table S3 and Table S4 summarize all descriptive and statistical values.

**Figure 3.**
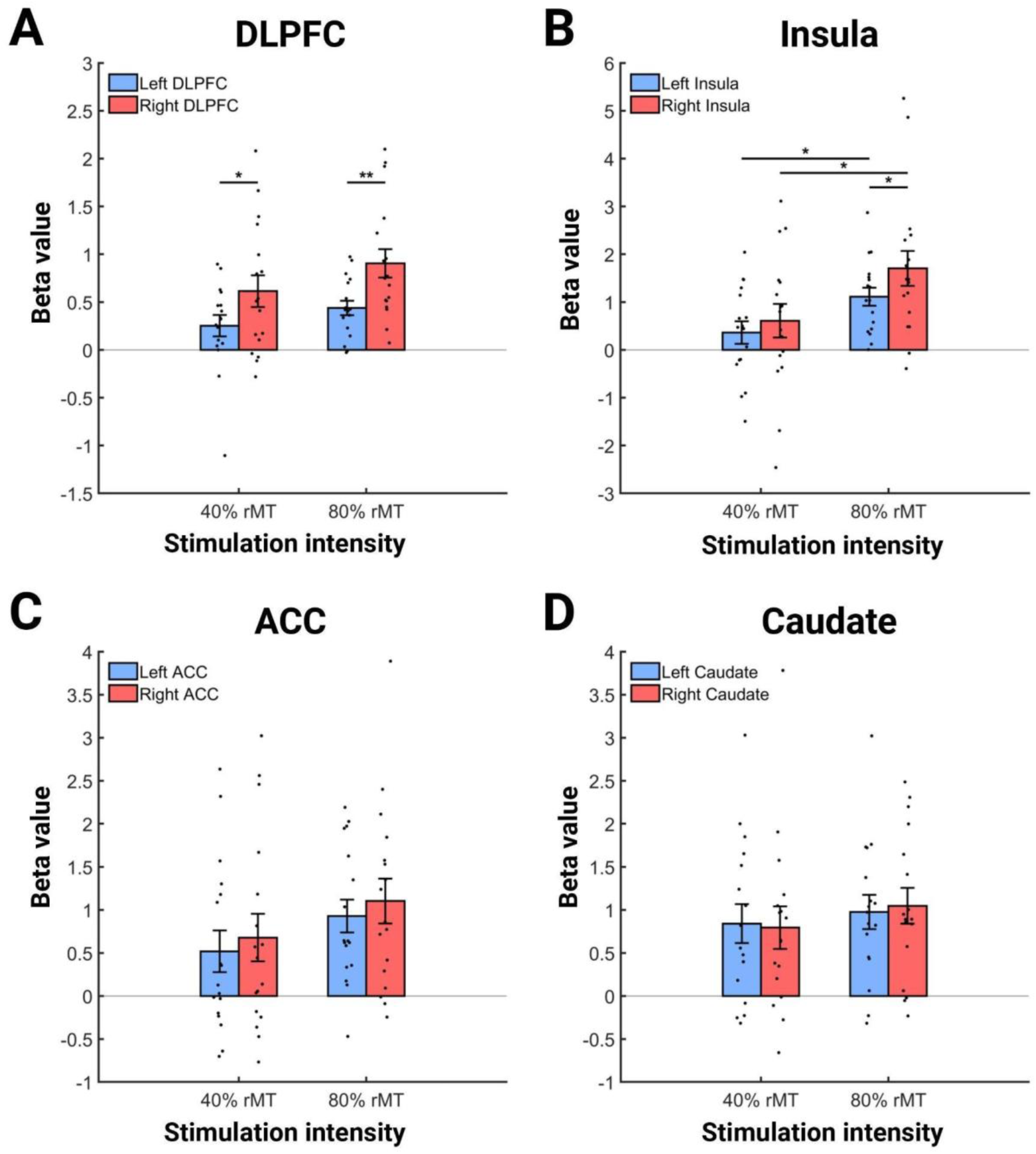
10 Hz rTMS-evoked BOLD responses in bilateral ROIs at 40% rMT and 80% rMT. DLPFC = dorsolateral prefrontal cortex; Insula = anterior insula; ACC = rostral anterior cingulate cortex; Caudate = dorsal caudate nucleus. Black dots represent individual subjects. Error bars show ± SEM. * p < .05; ** p < .01; *** p < .001.

### 3.3 Individual BOLD responses

Responses in the bilateral DLPFC extracted from the spherical ROIs were further analysed to highlight individual responses and contributions to the group-level effects. We found that BOLD responses were increased at higher stimulation intensity in 59% of the subjects (10/17) in the left DLPFC (Figure 4A) and in 71% of the subjects (12/17) in the right DLPFC (Figure 4B). Additionally, 76% of the subjects (13/17) exhibited a greater prefrontal BOLD response in the right DLPFC compared to the left DLPFC at both 40% rMT (Figure 4C) and 80% rMT (Figure 4D).

**Figure 4.**
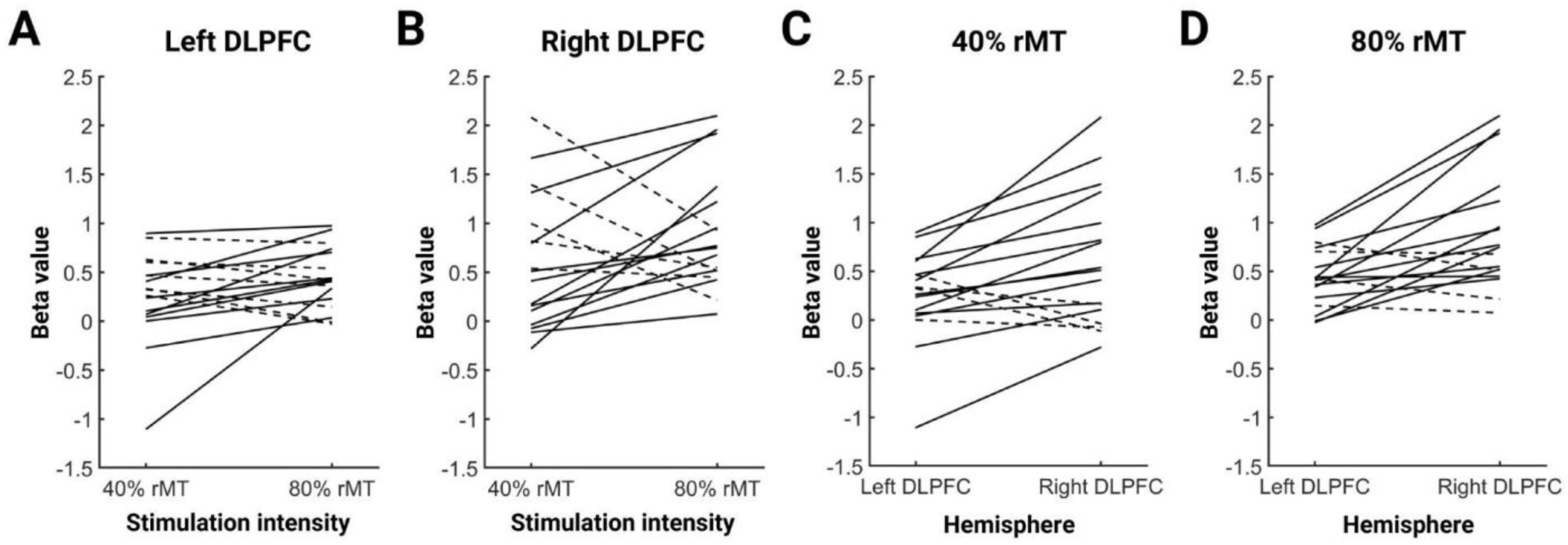
Individual responses to low and high 10 Hz rTMS in bilateral DLPFCs. (A) Dose-response relationship in the left DLPFC between 40% rMT and 80% rMT for each subject. (B) Dose-response relationship in the right DLPFC between 40% rMT and 80% rMT for each subject. The solid lines indicate individuals who showed a stronger activation with 80% rMT intensity, while the dashed lines indicate individuals who showed a stronger activation with 40% rMT intensity. (C) Hemispheric lateralization effect in DLPFC at 40% rMT for each subject. (D) Hemispheric lateralization effect in DLPFC at 80% rMT for each subject. The solid lines indicate individuals who showed a stronger activation in the right DLPFC, while the dashed lines indicate individuals who showed a stronger activation in the left DLPFC.

### 3.4 Temporal dynamics of BOLD response

To examine the time-dependent cumulative effects of consecutive rTMS trains on cortical responses, we divided the 10-minute stimulation period into three equal blocks of 3 minutes and 20 seconds (200 stimuli/block), aligning with the duration of a full iTBS protocol (15, 24). In the first block, BOLD activation matched the pattern seen across the entire stimulation period, with widespread activation in the bilateral DLPFC, cingulate cortex, precentral gyrus, insula, thalamus, and striatum (Figure 5). The second and third blocks showed a gradual decrease in whole-brain BOLD cluster activation. By the third block of 80% rMT stimulation, significant clusters were present only in the bilateral ACC, right insula, right inferior frontal gyrus, right superior parietal lobule, right thalamus, and right striatum (both caudate nucleus and putamen) (Figure 5B). The second and third block of 40% rMT were characterized by significant bilateral striatal activation (both putamen and right caudate) (Figure 5A). Similar effects were seen when dividing blocks into six blocks of 100 pulses (Figure S5).

**Figure 5.**
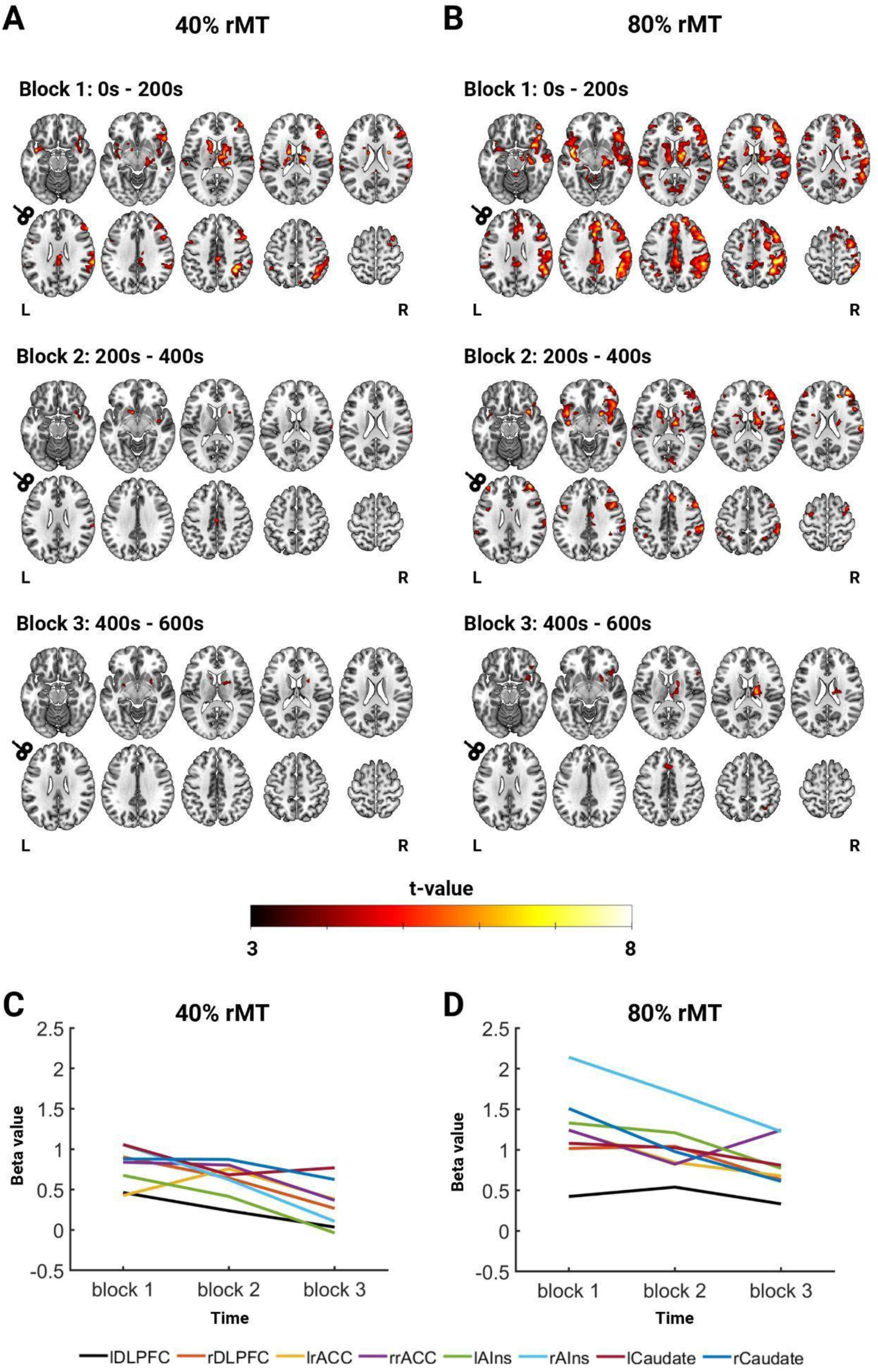
Temporal dynamics of BOLD response during 10 Hz rTMS. To examine how the BOLD response developed over time, the full 10 Hz rTMS protocol (10 minutes, 60 trains, 600 pulses) was divided into three blocks (each 3 minutes and 20 seconds, 20 trains, 200 pulses). (A) Whole brain activation maps across three blocks during 40% rMT. (B) Whole brain activation maps across three blocks during 80% rMT. All activation maps are thresholded at voxel-level p < .001 and cluster-level p < .05 FWE corrected. Axial slices have MNI Z-coordinates = -16 -8 12 18 22; 26 34 42 52 62. (D, E) Mean beta values in each ROI plotted for each block. lDLPFC = left dorsolateral prefrontal cortex; rDLPFC = right dorsolateral prefrontal cortex; lrACC = left rostral anterior cingulate cortex; rrACC = right rostral anterior cingulate cortex; lAIns = left anterior insula; rAIns = right anterior insula; lCaudate = left dorsal caudate nucleus; rCaudate = right dorsal caudate nucleus.

To rule out the possibility of visualization artifacts from the implicit baseline, we further analyzed the group-average ROI beta values across time windows (Figure 5C-D). Overall, BOLD activity initially plateaued, followed by a modest reduction over time. However, a more pronounced decrease in activation was observed in the right anterior insula and an increase in the right ACC at 80% rMT (Figure 5E).

## 4. Discussion

The present study aimed to characterize neural responses to a 10 Hz rTMS protocol over the left DLPFC in healthy individuals using interleaved TMS-fMRI. We found that 10 Hz rTMS elicited BOLD responses in prefrontal regions, ACC, insula, striatum, thalamus, as well as auditory and somatosensory areas. Notably, the 10 Hz rTMS effects in the DLPFC were lateralized towards the contralateral (right) DLPFC. Dosage-dependent responses in the DLPFC between 40% and 80% rMT exhibited substantial inter-individual variability. Compared to our previous interleaved TMS-fMRI study applying iTBS over the left DLPFC with identical pulse numbers and intensities (Chang et al., 2024b), the 10 Hz rTMS effects showed distinct patterns of activation.

### 4.1 rTMS-evoked brain activity patterns

Previous interleaved 10 Hz TMS-fMRI studies of prefrontal regions in healthy subjects typically used shorter 10 Hz bursts (i.e., 3-8 pulses) and fewer total pulses per experimental condition (Hawco et al., 2017; Caparelli et al., 2022; Tik et al., 2023a, 2023b). The 10 Hz rTMS protocol in the current study with 600 stimuli (1 s ON, 9 s OFF) represents the longest application of a prefrontal 10 Hz rTMS protocol inside a MR scanner to date and parallels a full 600-stimuli iTBS protocol (Huang et al., 2005; Chang et al., 2024b).

We found that 10 Hz rTMS modulated neural activity within the target DLPFC and in interconnected cortical and subcortical regions. The observed BOLD signal increases in regions including the ACC, insula, striatum, and thalamus may point to a modulation of the salience network, a potential depression biomarker (Downar et al., 2016; Seeley, 2019). Earlier TMS-fMRI work stimulating the left DLPFC during rest with either single-pulses or 10 Hz bursts reported similar activation patterns underneath the coil and in distant areas (Hanlon et al., 2013; Vink et al., 2018; Dowdle et al., 2018; Tik et al., 2023a, 2023b). Activation of salience network nodes may also be related to auditory and somatosensory responses to the TMS pulse, or anticipation and attentional effects (Iannetti & Mouraux, 2010; Dowdle et al., 2018).

It was evident that group-level effects were strongly driven by the first few minutes of the 10-minute stimulation protocol. Notably, local activity was observed only during the first block, while activation shifted to remote areas as the protocol progressed (Figure 5; Figure S4). Our findings point to robust fronto-striatal-thalamic network engagement during left DLPFC 10 Hz rTMS, as these were stable across all temporal windows for 80% rMT as well as 40% rMT. Previous research has identified strong structural and functional connections between the DLPFC and striatal areas (Leh et al., 2007; Di Martino et al., 2008). Moreover, rTMS over the DLPFC modulates subcortical dopaminergic neurotransmission in healthy subjects (Strafella et al., 2001) and in those with depression (Pogarell et al., 2006, 2007). Accordingly, the frontal-striatal-thalamic pathway may play a key role in the antidepressant effect of rTMS in depression (Avissar et al., 2017). In future studies, interleaved TMS-fMRI in clinical populations may be therefore used to elucidate whether fronto-striatal-thalamic network engagement predicts better clinical response to rTMS. Another main research question is to what extent such initial local and sustained remote effects during 10 Hz rTMS contribute to its therapeutic efficacy. A deeper understanding of both temporal dynamics and interindividual variance of effects may guide choices of stimulation patterns and durations required to achieve optimal neurophysiological effects and clinical outcomes.

### 4.2 High and low intensity of 10 Hz rTMS

The field of TMS lacks an established sham control capable of fully isolating the neurophysiological effects of TMS independently of accompanying somatosensory responses (Arana et al., 2008; Duecker & Sack, 2015; Bagali et al., 2023). In this study, we employed an active intensity control at 40% rMT, similar to our previous study (Chang et al., 2024b). This study comprised separate sessions for each intensity, whereas previous TMS-fMRI studies often compared multiple stimulation intensities within a single session (Nahas et al., 2001; Baudewig et al., 2001; Moisa et al., 2010; Tik et al., 2023a, 2023b), which may have led to carry-over effects and reduced signal to noise ratios (Seewoo et al., 2019).

In the left DLPFC target region, we did not observe a dose-dependent BOLD effect comparing 40% and 80% rMT intensity. While linear TMS dose-responses are consistently detected in M1 (Bohning et al., 1999; Hanawaka et al., 2009; Navarro de Lara et al., 2017; Chang et al., 2024b), findings in studies using interleaved TMS-fMRI over the left DLPFC are mixed regarding dose-dependent effects (Nahas et al., 2001; Dowdle et al., 2018; Tik et al., 2023b; Chang et al., 2024b). In our study, only 59% of the subjects showed increased left DLPFC activity with higher stimulation intensity, potentially preventing a group-level effect. Similarly, previous inconclusive findings may have been due to the large between-subject variability (Tik et al., 2023b).

Dose-response effects (40% vs. 80% rTMS) were exclusively observed in subcortical regions. Whole-brain analysis revealed a significant difference between the two stimulation intensities in the hippocampus. The DLPFC and hippocampus are directly connected through distinct functional pathways (Goldman-Rakic et al., 1984). It is suggested that remote areas rather than the stimulated DLPFC show compensatory patterns to rTMS, with reported linear activity increases in subcortical regions with increasing stimulation intensities of rTMS over left DLPFC (Tik et al., 2023b). The insula was also increasingly activated by higher intensity stimulation, potentially reflecting discomfort from TMS (Iannetti & Mouraux, 2010).

### 4.3 Comparing outcomes of 10 Hz rTMS and iTBS with interleaved TMS-fMRI

There was significant lateralization of effects during left DLPFC 10 Hz rTMS towards the contralateral hemisphere, i.e., the right DLPFC and right insula. This was evident in most subjects (Figure 4C-D), and persisted throughout the entire stimulation protocol (Figure 5, Figure S4). Previous TMS-fMRI (Tik et al., 2023b) and TMS-EEG studies (Ye et al., 2022; Ross et al., 2023) have similarly shown evidence that left prefrontal 10 Hz rTMS has its strongest effects in the right hemisphere. In TMS-fMRI studies, there are often no local but only distal BOLD effects (Bergmann et al., 2021; Rafiei & Rahnev, 2022), and the mechanisms underlying this observation are not clear. First, activation of distant regions could theoretically be due to nonspecific effects associated with TMS-fMRI, and it may be tempting to expect and interpret effects in proximal regions as more stimulation target related than effects in distal regions. Second, cortical stimulation specific effects could be explained by transsynaptic effects of target modulation, but it is not clear whether these distant effects are due to real excitatory stimulation (as often assumed) or mediated by interference with the complex inhibitory control of cortical microcircuits (Hamada et al., 2013).

Interestingly, the present findings largely differ from our previous findings of left DLPFC stimulation by a 600-stimuli iTBS protocol at 40% and 80% rMT, where the BOLD activation in the left DLPFC was significantly higher than in the right DLPFC (Chang et al., 2024b). Regarding their clinical applications, iTBS, which is a patterned high-frequency rTMS protocol delivering bursts of 50 Hz triplets repeated at 5 Hz (Huang et al., 2005), and 10 Hz rTMS, are often regarded as equivalent in terms of their therapeutic effects in MDD (Blumberger et al., 2018; Bulteau et al., 2022; Spitz et al., 2022). In direct comparison, however, the neurophysiological action of these two widely used stimulation protocols for MDD – 10 Hz rTMS and iTBS – remain largely unexplored, and their clinical equivalence has been recently challenged (Li et al., 2023; Wada et al., 2025). The findings of both interleaved TMS-fMRI studies together suggest that 10 Hz rTMS and iTBS are protocols with apparently distinct neurophysiological properties in terms of target engagement and propagation patterns. These protocols differed only in how the TMS pulses are grouped together over time, which means that the timing of TMS pulses could play a pivotal role in determining acute brain responses, possibly by recruiting distinct neuronal populations.

Early findings for M1 (Pascual-Leone et al., 1994; Wassermann et al., 1998; Huang et al., 2005) have led to the misguided assumption that rTMS protocols are inherently physiologically excitatory (when the stimulation frequency is higher than 5 Hz) or inhibitory (when the frequency is lower than 5 Hz) in their cortical effects, even when they are applied to non-motor cortical sites such as the DLPFC (Hussain & Freedberg, 2025). Indeed, neuromodulatory effects of standard rTMS in motor regions may be less robust than initially expected, despite the fact that these protocols were originally established for motor system plasticity induction. Recent studies investigating common variants of rTMS over M1 showed inter- and intra-individually highly variable effects with little consistency (Schilberg et al., 2017; Boucher et al., 2021; Magnuson et al., 2023).

While left DLPFC activation progressively increased over the 3-minute iTBS protocol (14), we found no such cumulative effect with 10 Hz rTMS. Strikingly, iTBS and 10 Hz rTMS produced similar activation patterns at the lower intensity: the higher frequency (or shorter duration) of iTBS may induce plasticity underneath the coil more effectively. Whether 10 Hz rTMS would require higher intensities or longer durations (such as in clinically used) (O’Reardon et al., 2007) to achieve similar cumulative effects remains speculative and warrants further research.

### 4.4 Limitations & Future directions

Simultaneous TMS and fMRI acquisition is technically challenging and inherently introduces rigid artifacts into the imaging data (Mizutani-Tiebel et al., 2022). In addition, prolonged rTMS inside the MR scanner (i.e., ∼10 minutes for 10 Hz rTMS) can cause substantial discomfort compared to offline rTMS, leading to increased head motion and lower signal-to-noise. While a larger sample size would potentially improve the robustness and reliability of the findings, the high costs associated with TMS-fMRI data acquisition need to be considered.

In this study, we modified the standard therapeutic 10 Hz rTMS protocol (100%-120% rMT; 4 s ON, 26s OFF; 3000 pulses; 37.5 minutes) (O’Reardon et al., 2007), implementing subthreshold intensities (40% and 80% rMT) and shorter duration (600 pulses, 10 minutes), for three primary reasons: 1) technical constraints of the TMS coil (e.g., heating, larger MR artifacts), 2) reducing participant discomfort, and 3) facilitating a direct comparison with the full iTBS protocol (i.e., 600 pulses/session and 80% MT), as it was also implemented in prior work by Chang et al. (2024b). Further advancements in interleaved TMS-fMRI technology ought to enable closer alignment with both offline and online therapeutic rTMS protocols in the future. Whereas the inclusion of more suprathreshold stimulation intensities (Tik et al., 2023b) or other stimulation targets, e.g., M1 (Chang et al., 2024b), would have been interesting to investigate the specificity of effects observed here, this was generally restricted by these experimental limitations.

Importantly, these findings in a healthy population may have limited translation value and applicability to clinical populations. rTMS may affect brain activity differently in individuals with pathological brain functions. Extending TMS-fMRI research to clinical populations with network disorders (e.g., MDD, schizophrenia, obsessive-compulsive disorder) is necessary to establish clinical relevance and determine its potential in identifying predictive biomarkers for neuromodulation-based treatment in psychiatry.

## 5. Conclusions

Using interleaved TMS-fMRI, this study provides evidence of distinct prefrontal target engagement and propagation patterns during left prefrontal 10 Hz rTMS. By comparing the current findings on 10 Hz rTMS to previous findings from our lab on iTBS, this leads us to speculate that distinct neurophysiological mechanisms may be involved in 10 Hz rTMS and iTBS that determine the propagation of stimulation effects to interconnected brain regions. Further research is necessary to elucidate the underlying mechanisms of these therapeutic rTMS protocols and to determine how acute effects captured with TMS-fMRI relate to treatment response in clinical populations. In sum, interleaved TMS-fMRI using full rTMS protocols represents a highly promising approach for investigating rTMS protocols in terms of their immediate effects on brain circuits including individual differences which could lead to the development of clinically effective and personalized rTMS in the future.

## Funding information

This project was funded by the Federal Ministry of Education and Research within the ERA-NET NEURON program (Grant No. FKZ: 01EW1903: DisCoVeR) and the German Center for Mental Health (Grant No. 01EE2303, project MUC6). The procurement of the Prisma 3T MRI scanner was supported by the Deutsche Forschungsgemeinschaft (Grant No. INST 86/1739-1 FUGG). LB’s work was part of the funding program of the Medical Faculty of LMU (FöFöLe Plus, Munich Clinician Scientist Program). MT’s travel costs and research stays were supported by the visiting scholarship artificial intelligence programme (BaCaTeC 2022/01, Machine learning-based optimization for personalized brain stimulation therapy). PT’s work was funded by the Deutsche Forschungsgemeinschaft (Grant No. TA 857/3-2). MC’s work was part of the funding program of the Medical Faculty of LMU (FöFöLe, Munich Clinician Scientist Program)

## Data and code availability statement

Access to the completed MRI data and unthresholded statistical maps are available upon request. The MRI can be obtained upon formalizing a data sharing agreement, though not publicly accessible due to challenges in achieving complete anonymization.

## CRediT authorship contribution statement

TvH: Conceptualization, Methodology, Formal analysis, Investigation, Data Curation, Writing - Original Draft, Writing - Review & Editing, Visualization, Project administration. K-YC: Conceptualization, Methodology, Validation, Investigation, Writing - Review & Editing, Supervision, Project administration. MT: Conceptualization, Methodology, Validation. PT: Writing - Review & Editing. JB: Writing - Review & Editing. LB: Writing - Review & Editing, Project administration. FP: Conceptualization, Methodology, Validation, Resources, Writing - Review & Editing, Supervision, Project administration, Funding acquisition. DK: Conceptualization, Methodology, Validation, Resources, Writing - Review & Editing, Supervision, Project administration, Funding acquisition. MC: Conceptualization, Methodology, Validation, Investigation, Writing - Review & Editing, Supervision, Project administration, Funding acquisition.

## Acknowledgements

We thank Yuki Mizutani-Tiebel and Jualian Melcher for assisting TMS-fMRI data collection. We thank Christian Windischberger (Center for Medical Physics and Biomedical Engineering at the Medical University of Vienna, Austria) for providing expertise knowledge and advice in concurrent TMS-MRI, as well as Patrik Kunz (Localite GmbH, Bonn, Germany) and Matthias Kienle (MagVenture A/S, Farum, Denmark) for their support with the technical setup. This work is a part of a research project by TvH conducted at LMU Klinikum Munich as part of his Master’s degree at the University of Amsterdam, The Netherlands.

## Disclosures

FP is a member of the European Scientific Advisory Board of BrainsWay Inc. (Jerusalem, Israel), and the International Scientific Advisory Board of Sooma (Helsinki, Finland); he has received speaker honoraria from Mag&More GmbH, the neuroCare Group (Munich, Germany), and BrainsWay Inc.; his lab has received support with equipment from neuroConn GmbH (Ilmenau, Germany), Mag&More GmbH, and BrainsWay Inc. TvH, K-YC, MT, PT, JB, LB, DK, & MC reported no potential conflicts of interest.

## Appendix. Supplementary Methods

### fMRI Preprocessing

Data preprocessing and statistical analysis were performed using MATLAB (Mathworks, Natrick, MA) and the SPM12 toolbox (Wellcome Trust Centre for Neuroimaging, London, United Kingdom), as well as ANFI (http://afni.nimh.nih.gov/afni) and ANTs (http://stnava.github.io/ANTs). Preprocessing of the fMRI data included the following steps: 1) segmentation and normalization of the anatomical images to Montreal Neurological Institute (MNI) standard space with deformation fields using CAT12 (http://www.neuro.uni-jena.de/cat/); 2) bias-field correction of the functional images with N4BiasFieldCorrection implemented in ANTs; 3) despiking of the functional images using ANFI; 4) realignment of the functional images (SPM12); 5) coregistration of the functional images to the anatomical images (SPM12); 6) normalization of coregistered functional images to MNI standard space based on deformation fields derived from anatomical normalization (SPM12); and 7) spatial smoothing using a 6 mm FNWH Gaussian kernel (SPM12). Subjects with mean framewise displacement greater than 0.3 mm were excluded from all further analysis (Power et al., 2012). Data quality was checked after each preprocessing step via visual inspection.

In order to reduce physiological noise (e.g., motion, cerebrospinal fluid (CSF) pulsations) and artifacts that may have been introduced by the interleaved TMS-fMRI setup (e.g, leakage currents, mechanical vibrations, RF interference due to the TMS hardware) (Bergmann et al., 2021; Riddle et al., 2022; Mizutani-Tiebel et al., 2022), an independent component analysis (ICA) was performed using FSL MELODIC (Jenkinson et al., 2012). Time signals were decomposed into 25 components and denoised via manual rejection (Griffanti et al., 2017). An average of 20.42 (SD = 1.52) components were removed across both conditions.

### E-field modeling

The open-source software package SimNIBS v4.1.0 was used for realistic calculations of the electric field (E-field) induced by TMS (Thielscher et al., 2015). High-resolution anatomical T1w and T2w images (TE = 383 ms, TR = 5000 ms, TA = 4:57 m, TI =1800 ms, flip angle = 8°, voxel size = 1.0 × 1.0 × 1.0 mm, number of slices = 176, slice thickness = 1 mm, FOV = 256 mm) from the baseline session were segmented into gray matter, white matter, and CSF, to construct realistic head models for each subject. Using stimulation intensity output values from the TMS stimulator (di/dt = A/µs), the individualized E-field on the cortex was simulated for each session. The location and orientation of the TMS coil were derived from the first markers recorded by the neuronavigation system. Three subjects were excluded in the 40% rMT (n = 14) condition and two subjects in the 80% rMT (n = 15) condition for E-field analysis because stimulation markers or intensity values were not properly recorded. Mean E-field strength in the left DLPFC was extracted using a spherical ROI (radius = 10 mm, x, y, z = -38, 44, 26) and correlated to mean ROI beta values in left DLPFC (Pearson’s correlation, p < .05).

The E-field induced by 10 Hz rTMS was focally located at the target site in the left DLPFC at 40% rMT (M = 28.12 V/m, SD = 6.85) and 80% rMT (M = 57.84 V/m, SD = 10.10) [paired t-test: t(12) = 14.662, p < .001] (Figure S3). However, there was a lack of statistically significant correlations between E-field strength and individual BOLD signal in the left DLPFC at both 40% rMT [r = -0.077, p = 0.793] and 80% rMT [r = 0.363, p = 0.184] (Figure S3).

## Appendix. Supplemental Tables

**Table S1.** Subject demographics and mean framewise displacement. Framewise displacement was calculated according to Power et al. (1)

| Subject ID | Age | Sex | rMT (% MSO) | Mean FD - 40%<br>rMT (mm) | Mean FD - 80%<br>rMT (mm) |
| --- | --- | --- | --- | --- | --- |
| sub001 | F | 24 | 78 | 0.10 | 0.18 |
| sub002 | M | 56 | 69 | 0.21 | 0.12 |
| sub003 | F | 24 | 74 | 0.10 | 0.11 |
| sub005 | M | 32 | 66 | 0.06 | 0.07 |
| sub007 | F | 31 | 72 | 0.10 | 0.18 |
| sub008 | F | 28 | 80 | 0.09 | 0.09 |
| sub009 | F | 21 | 71 | 0.10 | 0.12 |
| sub010 | F | 25 | 70 | 0.09 | 0.11 |
| sub011 | M | 33 | 75 | 0.09 | 0.10 |
| sub012 | F | 26 | 89 | 0.11 | 0.09 |
| sub014 | M | 31 | 71 | 0.09 | 0.12 |
| sub015 | F | 27 | 61 | 0.09 | 0.10 |
| sub016 | M | 23 | 70 | 0.17 | 0.15 |
| sub017 | F | 24 | 75 | 0.09 | 0.12 |
| sub018 | F | 28 | 68 | 0.09 | 0.12 |
| sub019 | F | 23 | 87 | 0.10 | 0.13 |
| sub020 | M | 23 | 90 | 0.15 | 0.12 |

**Table S2.** Location and spherical size of regions-of-interest (ROI).

| ROI | Left hemisphere<br>MNI coordinates<br>(x, y, z) | Right hemisphere<br>MNI coordinates<br>(x, y, z) | Radius<br>sphere (mm) | Reference |
| --- | --- | --- | --- | --- |
| DLPFC | -38, 44, 26 | 38, 44, 26 | 10 | (Blumberger et al., 2018) |
| Rostral ACC | -5, 34, 28 | 5, 34, 28 | 5 | (Zhou et al., 2016) |
| Anterior insula | -44, 13, 1 | 47, 14, 0 | 10 | (Krönke et al., 2020) |
| Dorsal caudate<br>nucleus | -13, 15, 9 | 13, 15, 9 | 5 | (Park et al., 2020; Peng<br>et al., 2022) |

**Table S3.**
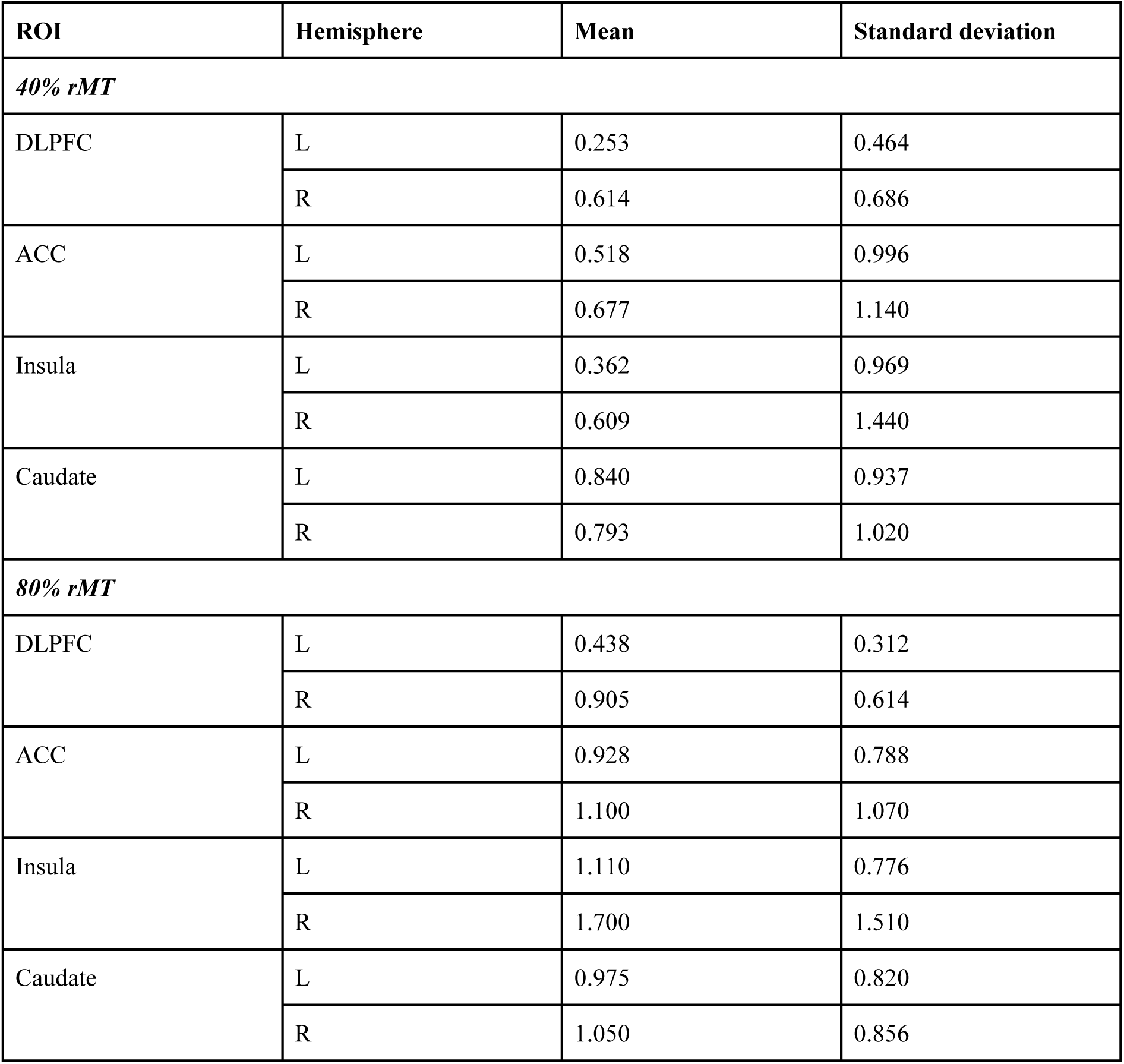
10 Hz rTMS descriptive statistics for each ROI.

**Table S4.** 10 Hz rTMS test statistics for each ROI.

|  | Test statistic | p-uncorrected | p-corrected | Significance |
| --- | --- | --- | --- | --- |
| ***DLPFC*** |  |  |  |  |
| Main effect: intensity | $F(1,16) = 3.194$ | 0.093 | - | - |
| Main effect: hemisphere | $F(1,16) = 15.337$ | 0.001 | - | * |
| Interaction effect | $F(1,16) = 0.659$ | 0.429 | - | - |
| left 40% rMT - left 80% rMT | $t(16) = -1.76$ | 0.098 | 0.196 | - |
| right 40% rMT - right 80% rMT | $t(16) = -1.60$ | 0.128 | 0.196 | - |
| left 40% rMT - right 40% rMT | $t(16) = -2.98$ | 0.009 | 0.027 | * |
| left 80% rMT - right 80% rMT | $t(16) = -3.67$ | 0.002 | 0.008 | ** |
| ***ACC*** |  |  |  |  |
| Main effect: intensity | $F(1,16) = 3.653$ | 0.074 | - | - |
| Main effect: hemisphere | $F(1,16) = 1.340$ | 0.264 | - | - |
| Interaction effect | $F(1,16) = 0.007$ | 0.932 | - | - |
| left 40% rMT - left 80% rMT | $t(16) = -1.92$ | 0.073 | 0.292 | - |
| right 40% rMT - right 80% rMT | $t(16) = -1.65$ | 0.118 | 0.354 | - |
| left 40% rMT - right 40% rMT | $t(16) = -1.03$ | 0.320 | 0.640 | - |
| left 80% rMT - right 80% rMT | $t(16) = -0.947$ | 0.358 | 0.640 | - |
| ***Insula*** |  |  |  |  |
| Main effect: intensity | $F(1,16) = 10.182$ | 0.006 | - | * |
| Main effect: hemisphere | $F(1,16) = 6.518$ | 0.021 | - | * |
| Interaction effect | $F(1,16) = 1.466$ | 0.244 | - | - |
| left 40% rMT - left 80% rMT | $t(16) = -3.01$ | 0.008 | 0.032 | * |
| right 40% rMT - right 80% rMT | $t(16) = -2.87$ | 0.011 | 0.033 | * |
| left 40% rMT - right 40% rMT | $t(16) = -1.26$ | 0.226 | 0.226 | - |
| left 80% rMT - right 80% rMT | $t(16) = -2.50$ | 0.024 | 0.048 | * |
| ***Caudate*** |  |  |  |  |
| Main effect: intensity | $F(1,16) = 0.735$ | 0.404 | - | - |
| Main effect: hemisphere | $F(1,16) = 0.018$ | 0.895 | - | - |
| Interaction effect | $F(1,16) = 0.753$ | 0.398 | - | - |
| left 40% rMT - left 80% rMT | t(16) = -0.636 | 0.534 | 1 | - |
| right 40% rMT - right 80% rMT | t(16) = -0.979 | 0.342 | 1 | - |
| left 40% rMT - right 40% rMT | t(16) = 0.390 | 0.702 | 1 | - |
| left 80% rMT - right 80% rMT | t(16) = -0.642 | 0.530 | 1 | - |

*Note.* P-values are FWE corrected. \* $p < .05$ ; \*\* $p < .01$ ; \*\*\* $p < .001$

## Appendix. Supplemental Figures

**Figure S1.**
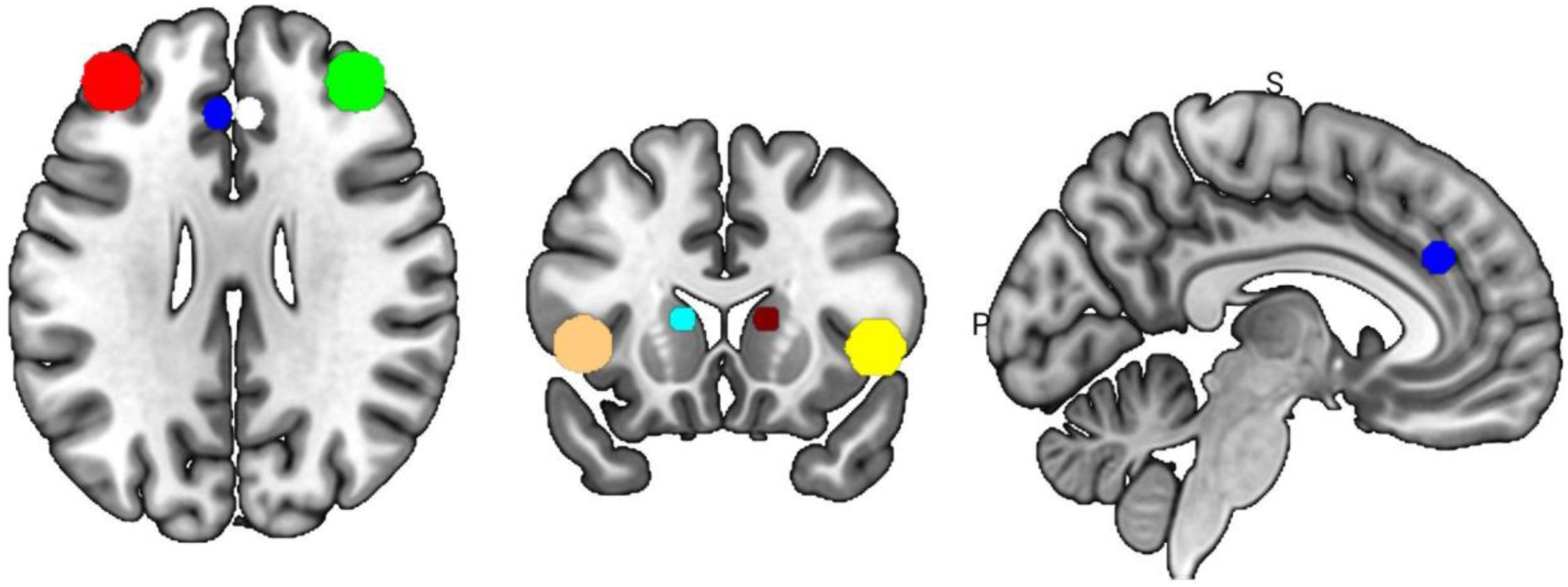
Location and spherical size of regions-of-interest (ROI). A spherical ROI with a 10 mm radius was adopted for bilateral DLPFC, as in previous publications from our lab (Chang et al., 2024a, 2024b). Other ROIs had deviating spherical sizes to prevent anatomical overlap. Red = left DLPFC; Green = right DLPFC; Navy = left rostral ACC; White = right rostral ACC; Orange = left anterior insula; Yellow = right anterior insula; Turquoise = left dorsal caudate nucleus; Bordeaux = right dorsal caudate nucleus.

**Figure S2.**
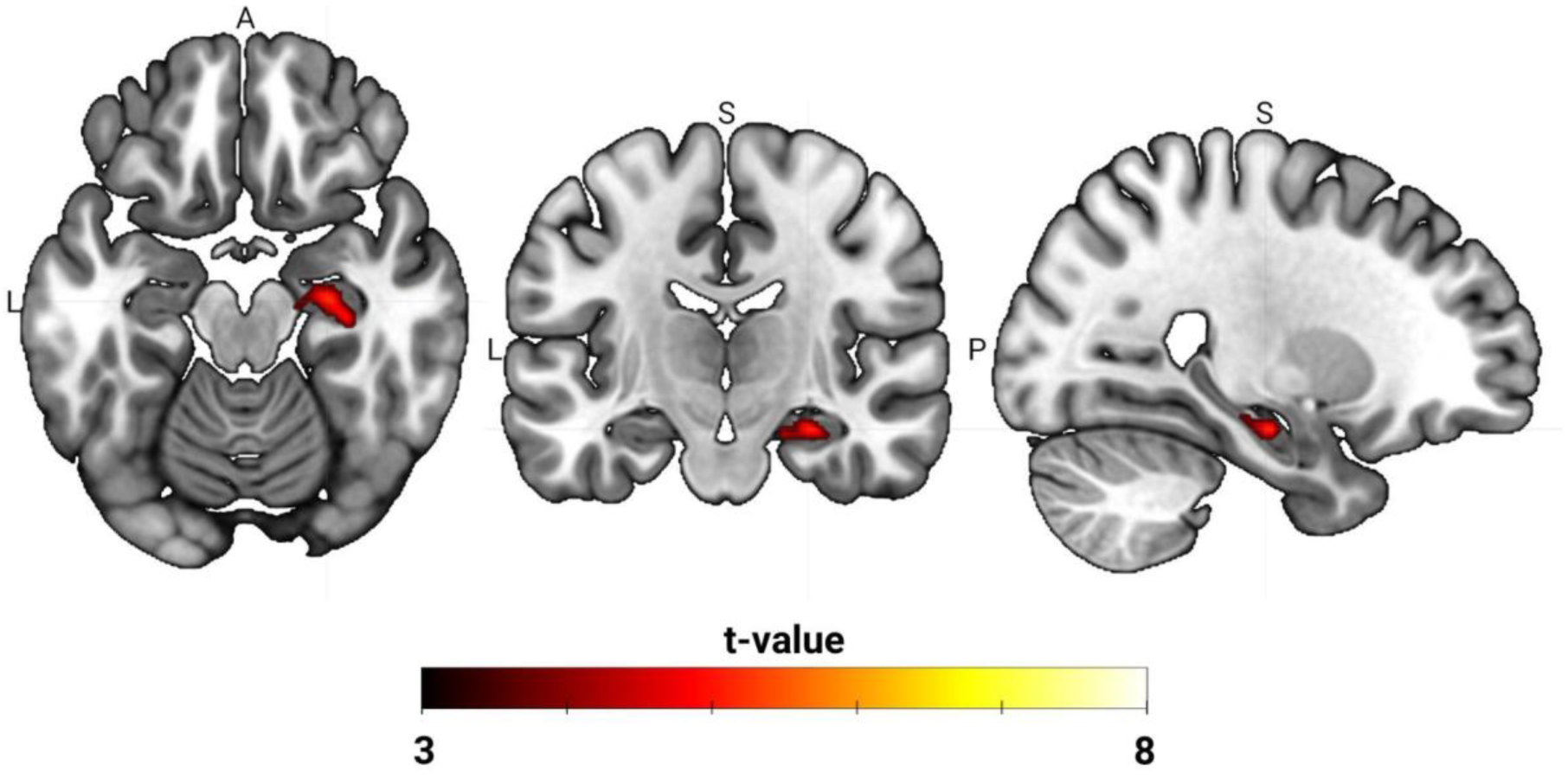
Contrast of 10 Hz rTMS-evoked BOLD responses 80% rMT > 40% rMT. A direct comparison of activation maps between 40% rMT and 80% rMT with paired t-test showed a cluster of significant difference in the right hippocampus (x, y, z = 26, -16, -18; k = 102; t = 5.31; Z = 3.98; Figure S2). Activation maps were thresholded at voxel-level p < .001 and cluster-level p < .05 FWE corrected. A significant cluster is shown in the right hippocampus (x, y, z = 26, -16, -18; k = 102; t = 5.31; Z = 3.98).

**Figure S3.**
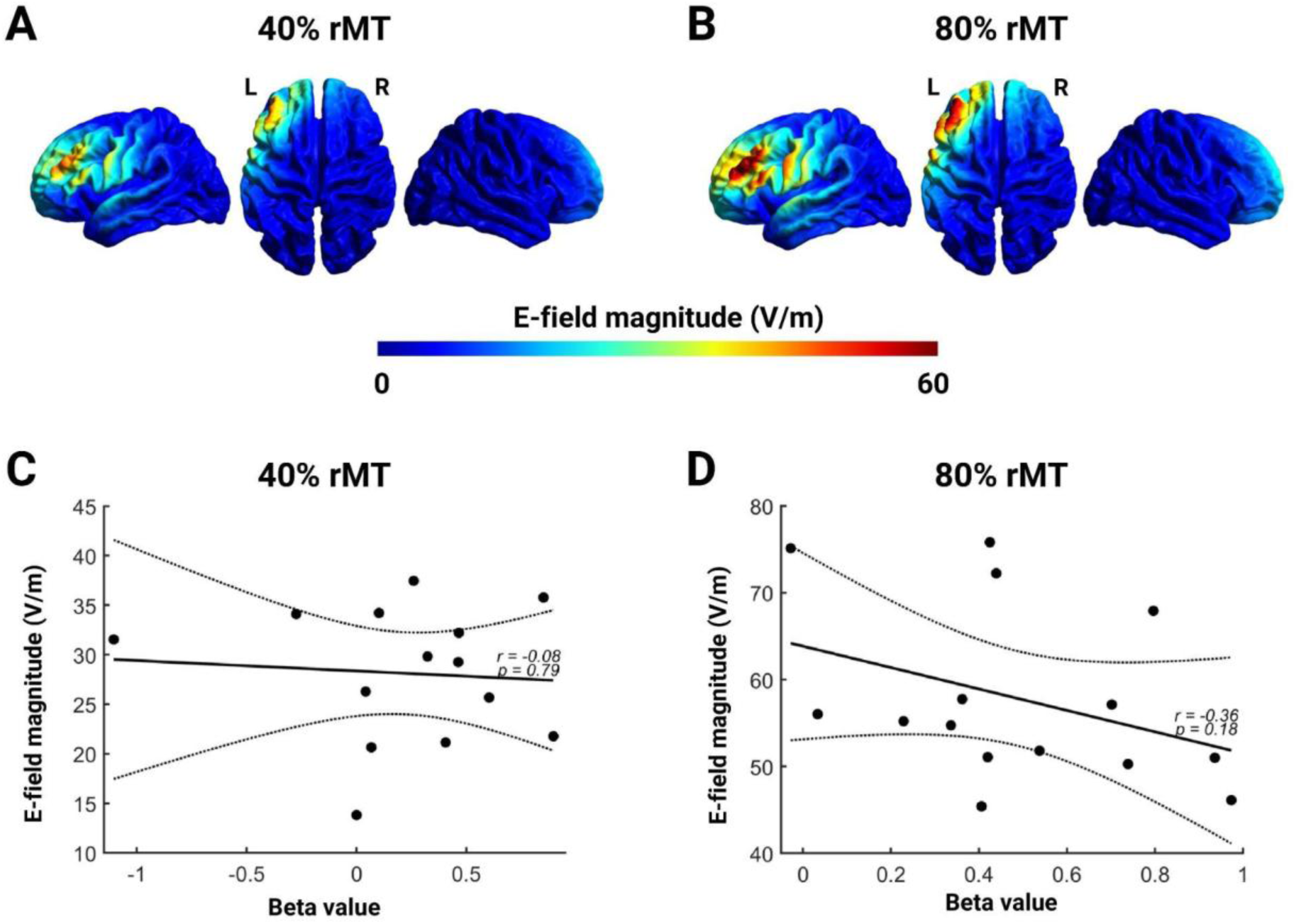
E-field simulations of 10 Hz rTMS over the left DLPFC. Mean E-Field strength showed specificity to the targeted location in the left DLPFC at 40% rMT (A) and 80% rMT (B). Individual beta values in the left DLPFC did not correlate with E-field strength at 40% rMT (C) or 80% rMT (D).

**Figure S4.**
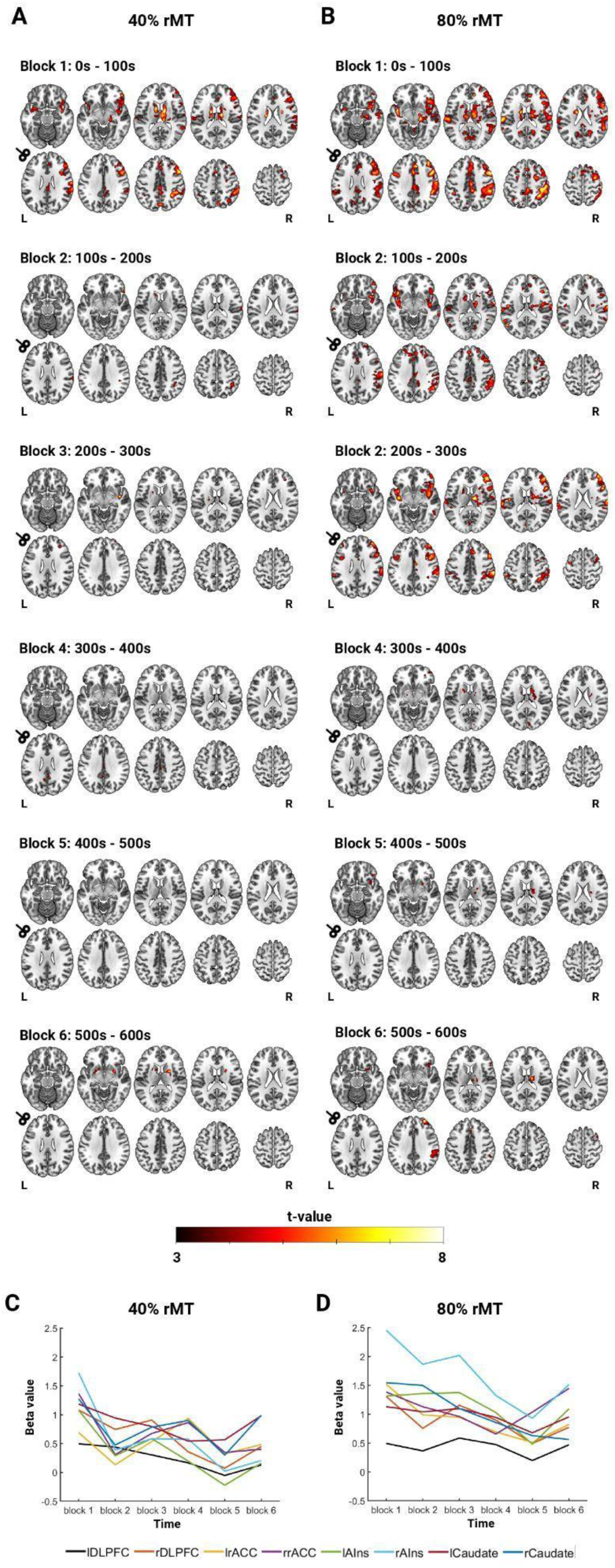
Temporal dynamics of BOLD response during 10 Hz rTMS protocol. The 10 Hz rTMS protocol lasted 10 minutes and consisted of 60 trains of 10 Hz rTMS (600 pulses total). To examine how the BOLD response developed over time, the full protocol was divided into six blocks, each containing 10 trains (100 pulses per block). Axial slices Z = -16 -8 12 18 22; 26 34 42 52 62.

**Figure S5.**
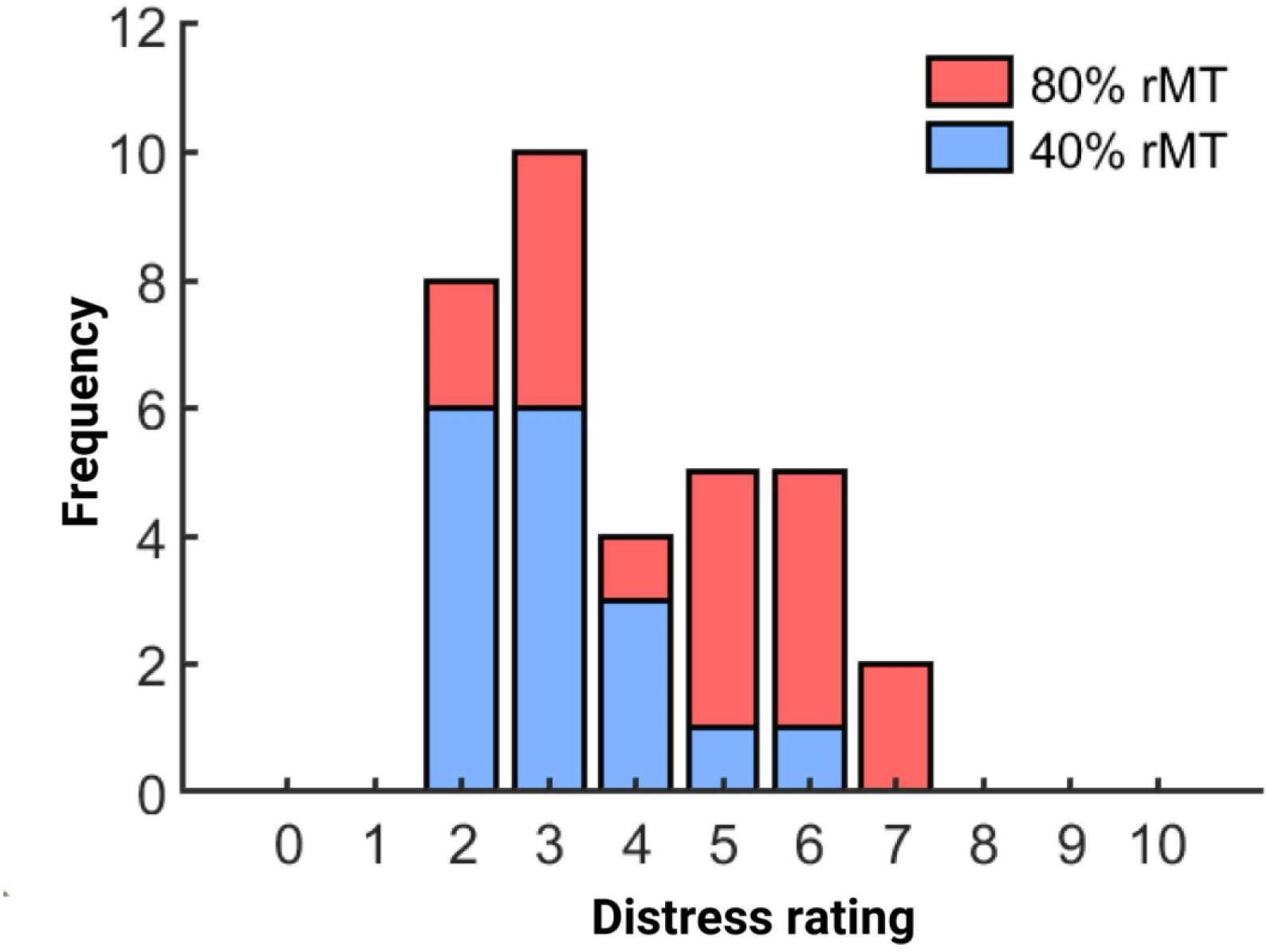
Distress ratings. Distress ratings, on a scale of 0 (no stress/pain)10 (extreme stress/pain), were recorded immediately after each stimulation session by asking participants to rate their pain and stress levels during the interleaved TMS-fMRI session. No subjects verbally reported experiencing pain during the experiment. (A) Distribution of distress ratings for both stimulation intensities. Distress ratings were higher for 80% rMT (M = 4.59, SD = 1.61) than for 40% rMT (M = 3.12, SD = 1.13). (B) Distress ratings sorted by session order.

## Appendix. Supplementary References

## Notes

### Competing Interest Statement

The authors have declared no competing interest.

